# Comparison of enrichment methods for efficient nitrogen fixation on a biocathode

**DOI:** 10.1101/2023.03.02.530809

**Authors:** Axel Rous, Gaëlle Santa-Catalina, Elie Desmond-Le Quémener, Eric Trably, Nicolas Bernet

## Abstract

The production of nitrogen fertilizers in modern agriculture is mostly based on the Haber-Bosch process, representing nearly 2% of the total energy consumed in the world. Low-energy bioelectrochemical fixation of N_2_ to microbial biomass was previously observed but the mechanisms of microbial interactions in N_2_-fixing electroactive biofilms are still poorly understood. The present study aims to develop a new method of enrichment of autotrophic and diazotrophic bacteria from soil samples with a better electron source availability than using H_2_ alone. The enrichment method was based on a multi-stage procedure. The first enrichment step was specifically designed for the selection of N_2_-fixing bacteria from soil samples with organic C as electron and carbon source. Then, a polarized cathode was used for the enrichment of autotrophic bacteria using H_2_ (hydrogenotrophic) or the cathode as electron source. This enrichment was compared with an enrichment culture of pure diazotrophic hydrogenotrophic bacteria without the use of a microbial electrochemical system. Interestingly, both methods showed comparable results for N_2_ fixation rates at day 340 of the enrichment with an estimated average of approximately 0.2 mgN_fixed_/L.d. Current densities up to −15 A/m^2^ were observed in the polarized cathode enrichments and a significant increase of the microbial biomass on the cathode was shown between 132 and 214 days of enrichment.These results confirmed an enrichment in autotrophic and diazotrophic bacteria on the polarized cathode. It was hypothesied that autotrophic bacteria were able to use either the H_2_ produced at the cathode or directly the cathode through direct electron transfer (DET) as more biomass was produced than with H_2_ alone. Finally, the analysis of the enriched communities suggested that *Desulforamulus ruminis* mediated microbial interactions between autotrophic anaerobic and heterotrophic aerobic bacteria in polarized cathode enrichment. These interactions could play a key role in the development of biomass in these systems and on N_2_ fixation. Based on these findings, a conceptual model on the functioning of mixed cultures N_2_-fixing electroactive biofilms was proposed.

## Introduction

Nitrogen is one of the essential elements for the growth of all living organisms, especially for cellular protein synthesis. In modern agriculture, ammonia is often used as a nitrogen source for plants (Bagali, 2012; Burris & Roberts, 1993; Masclaux-Daubresse et al., 2010). This compound is produced at industrial scale by the Haber-Bosch process which allows the reduction of N_2_ to NH_3_ at the expanse of large quantities of H_2_ and energy (Kandemir et al., 2013; Martín et al., 2019). This process is associated with significant CO_2_ emissions, due to the source of H_2_ obtained either by methane steam reforming or coal gasification (Martín et al., 2019). Alternatives for green H_2_ production, such as water electrolysis, are therefore nowadays considered to feed the Haber-Bosch process that is also contributing to a high energy demand of the process (Cherkasov et al., 2015). A more direct alternative is to reduce N_2_ directly on a cathode by chemical catalysts. However, these catalysts are not renewable and are currently not sufficiently selective regarding the hydrogen evolution reaction at ambient conditions (Deng et al., 2018; A. Liu et al., 2020).

Recently some authors proposed to use N_2_-fixing bacteria in association with electrochemical systems for N_2_ reduction at low energy cost (C. Liu et al., 2017; Rago et al., 2019). This idea was inspired by previous observations of microbial CO_2_ fixation on microbial cathodes in a process known as microbial electrosynthesis (A. Liu et al., 2020; Logan et al., 2019). The microbial fixation of both N_2_ and CO_2_ with a polarized cathode was demonstrated in two recent studies (C. Liu et al., 2017; Rago et al., 2019). First, Liu *et al*. (2017) demonstrated the growth of *Xanthobacter autotrophicus* in a hybrid organic-inorganic electrochemical system in the absence of nitrogen sources other than N_2_. Then, Rago et al. (2019) demonstrated N_2_ fixation through microbial electrosynthesis (MES) with a mixed microbial community (C. Liu et al., 2017; Rago et al., 2019).

Other works investigated the mechanisms with pure bacterial strain like Soundararajan et al. (2019) and Chen et al. (2020) with *Rhodopseudomonas palustris* and *Pseudomonas stutzeri*(Chen et al., 2020; Soundararajan et al., 2019). Yadav et al. (2022) demonstrated the possible use of N_2_-fixing bacteria as nitrogen source in a microbial electrosynthesis process of acetate(Yadav et al., 2022). Zhang et al. (2022) also worked on a system of N_2_ fixation in microbial electrolysis cell (MEC)(Zhang et al., 2022). In these works, the authors investigated the interactions existing between CO_2_ and N_2_ fixation microbial processes. Coupling capabilities of N_2_ fixation in bioelectrochemical systems as possible source of nitrogen for other biological systems was subject of interest. Indeed, such coupling can lead to the production of molecules of interest such as acetate by reducing the environmental impact of the use of reactive nitrogen often in the form of NH_4_Cl (Yadav et al., 2022). The work of Li et al. (2022) in a single-chamber system and highlighted an importance of synergy within an N_2_-fixing community in a microbial bioremediation system (Li et al., 2022).

All this work has demonstrated that it is possible to use a cathode as an electron source for biomass growth by fixing N_2_ and CO_2_. This biomass could then be used as a fertilizer with a low impact on the environment (Chakraborty & Akhtar, 2021; Rago et al., 2019; y. Hafeez et al., 2006). Although the proof of concept was made for this process, the microbial interactions supporting N_2_ fixation in this microbial electrochemical systems are still poorly understood. Different N_2_ fixation scenarios are indeed possible, such as: (i) fixation by a single population capable of fixing N_2_ and CO_2_ using the electrode as sole electron source, (ii) fixation by heterotrophic diazotrophic bacteria that can utilize the organic carbon produced by electro-autotrophic bacteria, (iii) fixation through an interaction between methanogenic archaea and methanotrophs that could use CH_4_ as an energy source for N_2_ fixation and (iv) fixation through an interaction mediated by direct interspecies electron transfer (DIET) between electro-autotrophic bacteria and diazotrophic bacteria (Rago et al., 2019). A better understanding of these interactions is essential to optimize N_2_ fixation in microbial electrochemical system.

In nature, biological N_2_ fixation is a key mechanism of the nitrogen cycle where atmospheric nitrogen is uptaken by living organisms (Bagali, 2012). It is carried out by so-called diazotrophic bacteria responsible for the transformation of N_2_ into NH_3_ (Kim & Rees, 1994). Some of these bacteria can be found on the roots of plants where they are living in symbiosis (Burris & Roberts, 1993; Franche et al., 2009). These microorganisms are able to fix N_2_ from the air, making it assimilable in the form of NH_4_^+^ or amino acids (L-glutamine, L-glutamate) which are further used in plants for protein or DNA synthesis (Burris & Roberts, 1993). In exchange, the bacteria use the organic matter of root exudates produced by the plants as carbon and energy sources. Among the diazotrophic bacteria, the genera *Frankia* and *Rhizobium spp*. are often associated with leguminous plant roots (Burris & Roberts, 1993; Peoples & Craswell, 1992). In contrast, other free-living N_2_-fixing bacteria such as *Azospirillum*, are able to fix N_2_ with or without interacting with plants and can use organic or inorganic materials to produce their own energy (Tilak et al., 1986).

In order to better understand the microbial mechanisms supporting N_2_ fixation in polarized cathode enrichment and produce biomass, this work aims at developing an enrichment method of cathodic biofilms for direct fixation of CO_2_ and N_2_. Rago et al. (2019) demonstrated the capacity of producing biomass from air, CO_2_ and a solid electrode polarized negatively (Rago et al., 2019). Here a specific enrichment of microorganisms capable of N_2_ fixation was developed. It was hypothesized that a multi-step enrichment with specific medium could select a microbial community able to fix N_2_ and CO_2_ supported by electrons brought by a cathode. It was assumed that these communities enriched by this procedure could use a large number of interactions leading to N_2_ fixation and biomass growth. For that, it was assumed that the enrichment in autotrophic diazotrophic bacteria in the presence of a cathode with pre-enrichment steps in presence of several electron donors (organic C and cathode) would be more efficient than enrichment cultures of hydrogenotrophic and diazotrophic bacteria. For this, soil samples were used as sources of N_2_-fixing bacteria, and successive enrichments in autotrophic bacteria in polarized cathode enrichment (PCE) were performed to select a electroactive biofilm capable of fixing N_2_ and CO_2_ with a cathode as sole electron source. The enriched biofilm was compared with a classical enrichment of N_2_-fixing hydrogen-oxidizing bacteria (HOB) in flasks (named H_2_ enrichment, H_2_E) (X. Hu et al., 2020; C. Liu et al., 2017).

## Methods

### Inoculum

Soil samples from a forest, agricultural crop field and a commercial compost were used as microbial inoculum sources. Samples were collected from forest and agricultural soils in the Haute Vallée de l’Aude, France. These sources were selected based on their assumed abundance of N_2_-fixing bacteria and their theoretical C/N ratio (Khan et al., 2016). One to two mg of each samples were used as inoculum in 50mL of medium for preliminary enrichment culture.

### Culture media

The culture media were both formulated on the basis of H3 medium (81 DSMZ) used for enrichment of soil autotrophic bacteria. The medium consisted of 2.3g KH_2_PO_4_ and 2.9g Na_2_HPO_4_ 2H_2_O per liter as buffer, 0.5g MgSO_4_ 7H_2_O, 0.01g CaCl_2_ 2H_2_O, 0.005g MnCl_2_ 4H_2_O, 0.005g NaVO_3_ H_2_O, and 5mL of SL-6 trace element solution per liter of medium, with 5mL of vitamin solution. The SL-6 trace element solution consisted of 0.1g ZnSO_4_ 7H_2_O, 0.03g MnCl_2_ 4H_2_O, 0.3g H_3_BO_3_ 0.2 CoCl_2_ 6H_2_O, 0.01g CuCl_2_ 2H_2_O, 0.02g NiCl_2_ 6H_2_O, and 0.03g Na_2_MoO_4_ 2H_2_O per liter of solution. The vitamine solution consisted of 10mg Riboflavin, 50mg Thiamine-HCl 2H_2_O, 50mg Nicotinic acid, 50mg Pyridoxine-HCl, 50mg Ca-Pantothenate, 0.1mg Biotin, 0.2mg Folic acid and 1mg Vitamin B_12_ for 100 mL of distilled water. Iron citrate was added to the enrichment bottles at a concentration of 0.05 g/L but not to the microbial bioelectrochemical systems in which the cathode was used as sole electron source when only CO_2_ was suplied. An organic carbon solution (organic C) was composed of 2g/L D-glucose, 1g/L yeast extract, 1g/L Na-acetate, 1g/L DL-malic acid, 1g/L Na-lactate, 1g/L Na-pyruvate, and 1g/L D-mannitol and used when indicated. NH_4_Cl was added at 1 g/L only when indicated. All enrichment procedures were maintained at 30°C and the pH was adjusted to 6.8 with NaHCO_3_ in the microbial bioelectrochemical systems and the enrichment cultures without organic C addition. When the organic C solution was used, the pH was adjusted between 6.3 and 6.5.

### Design of microbial electrochemical system

The electrochemical system used for our enrichment were composed of two chambers separated by an anion exchange membrane (AEM) (fumasep ® FAB-PK-130). The AEM was used to avoid the migration of NH_4_^+^ ions to the anodic chamber. Each chamber had a total volume of one liter. The pH of the reactors was adjusted to 6.8 at the beginning of each batch experiment. Each system had a 25 cm^2^ square carbon felt working electrode with a thickness of 0.7 cm and a 16 cm^2^ square Pt-Ir grid as a counter electrode. The carbon felt electrodes were conditioned using chemical treatment with HCl, a flush with ethanol and a heat treatment at +400°C as described elsewhere by Paul et al (Paul et al., 2018). The systems were inoculated with the flask enrichments used for N_2_ fixation in presence of organic C (N-free medium). After inoculation, the organic C supply was reduced to 10% of the initial supply in all reactors. The organic C supply was then totally replaced by CO_2_ supply after 60 days. Two of the systems were polarized and two other reactors were used as controls with open current voltage (non-polarized cathode enrichment, nPCE). The working electrodes of the polarized cathode enrichment (PCE) were poised at a potential of −0.940 V vs. saturated calomel electrode (SCE) used as a reference. The system was connected to a VMP3.0 potentiostat (BioLogic). The current was measured over time by a chronoamperometry method. The current intensity was used to monitor the availability of electrons in the electroactive biofilm. An increase of the current intensity was representative of an increase of the reduction reactions at the cathode. The increase of these reduction reactions, abiotic or not, was assumed to bring more electrons to the bacterial community. It was therefore assumed that an increase in current intensity was related to the enrichment of bacteria able to use the cathode as electron source (Zaybak et al., 2013). The current density *J* was calculated using the surface of the working electrode, i.e. 25 cm^2^.

Before inoculation of the reactors, an initial chronoamperometry measurement was performed along the first four days of operation with only organic carbon in the medium to determine the basal current density in absence of bacteria. Two other abiotic reactors for 15 days were then implemented to measure the current density in a medium without organic C. In order to validate the role of the cathode as electron source, two OCV (nPCE) reactors were carried out. The current densities of the abiotics reactors over a short period were therefore used as a reference to be compared with the measurements made after inoculation and monitor the increase of the activity of reduction at the cathode.

Electrons required for the production of microbial metabolites and for biomass growth was then used to calculate the Coulombic efficiency of the PCE according to the equations 1-5:

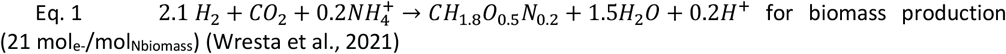

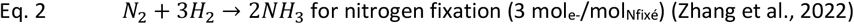

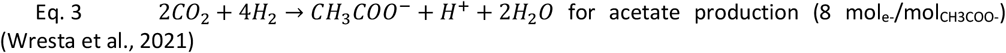

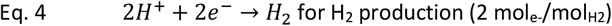

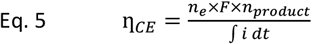

With η_*CE*_ the Coulombic efficiency in percentage of electron recovery in products, *n*_*e*_ moles of electrons per moles of product (mol_e-_/mol_product_) calculated from the stoichiometric equations, *F* the Faraday constant (96485 C.mol^-1^), *n*_*product*_ the number of mol_product_ and *i* the current intensity.

### Enrichment procedures

Two enrichment procedures based on a sequential procedure by enriching separately N_2_ fixation and the use of an inorganic electron source were carried out. Each soil sample (Forest, Leguminous and Compost) and a mix of all were used as inoculum in one batch of pre-enrichment. Pre-enrichment cultures were then carried out in the same medium as first enrichment step for each procedure (i.e. supplied with organic C for the first procedure and inorganic medium for the second) and for 20 days prior to be used as inoculum in the enrichment procedures. Samples of the pre-enrichment were considered as initial microbial community of the enrichments. Thus, at the end of pre-enrichment, the mixed culture was used to inoculate two bottles of each enrichment and all unmixed soil samples were mixed to inoculate four bottles of each enrichment.

<u>The first enrichment cultures, polarized cathode and non-polarized cathode enrichments (PCE and nPCE)</u> were performed in three steps:

- The first step was performed in a 120 mL bottle with N-free medium supplemented with organic C source. This first step was used to select N_2_-fixing bacteria using organic compounds as electron donors. The headspace was composed of an Ar/O_2_/N_2_ mixture (80/5/15) at 0.5 bar (absolute pressure). Cultures were carried out in bottles containing 50 mL of liquid and 70 mL of headspace. Subcultures of these enrichments were performed every 7 days for 6 weeks. The time between 2 subcultures was then reduced to 3-4 days using 10% of the volume of the previous culture (5mL/50mL).
- After 55 days of enrichment in 10 successive batches, the N_2_-fixing bacteria enriched cultures were used to inoculate the cathodic chambers of the polarized microbial electrochemical system (Polarized Cathode Enrichment, PCE) and the non-polarized microbial electrochemical system (non-Polarized Cathode Enrichment nPCE). The same inorganic medium supplemented with 10% of the organic C source was fed each week to start the enrichment of autotrophic bacteria. Inoculation of the cathode in presence of organic C sources was made to favor bacterial growth. 80% of the medium was renewed every second week to promote biofilm growth on the cathode. Headspace composition was monitored by GC and flushed with N_2_ if pO_2_ exceeded 10% of the gas volume. Organic C supply was stopped when a significant current density was measured in the polarized systems with regard to the controls.
- In the third and final enrichment step, CO_2_ was used as sole carbon source as presented in Figure 1. Organic C sources were then removed and the cathode was used as sole electron donor in reactor. Here, only bacteria able to use cathode as electron source by direct interaction or indirect with H_2_ were able to grow. A 80/20 (v:v) CO_2_/N_2_ atmosphere was set up in the headspace with trace amounts of O_2_ (< 5%). The medium was replaced every 15-30 days. The gas recycling vessel was filled with CO_2_ and was replaced with a new one when O_2_ exceeded 5% of the volume due to gas volume depletion. The nPCE controls were operated in the same conditions as the polarized systems but without monitoring the current density. The only available electron source in the nPCE controls was the organic C fed at the beginning of the enrichment.

**Figure 1.**
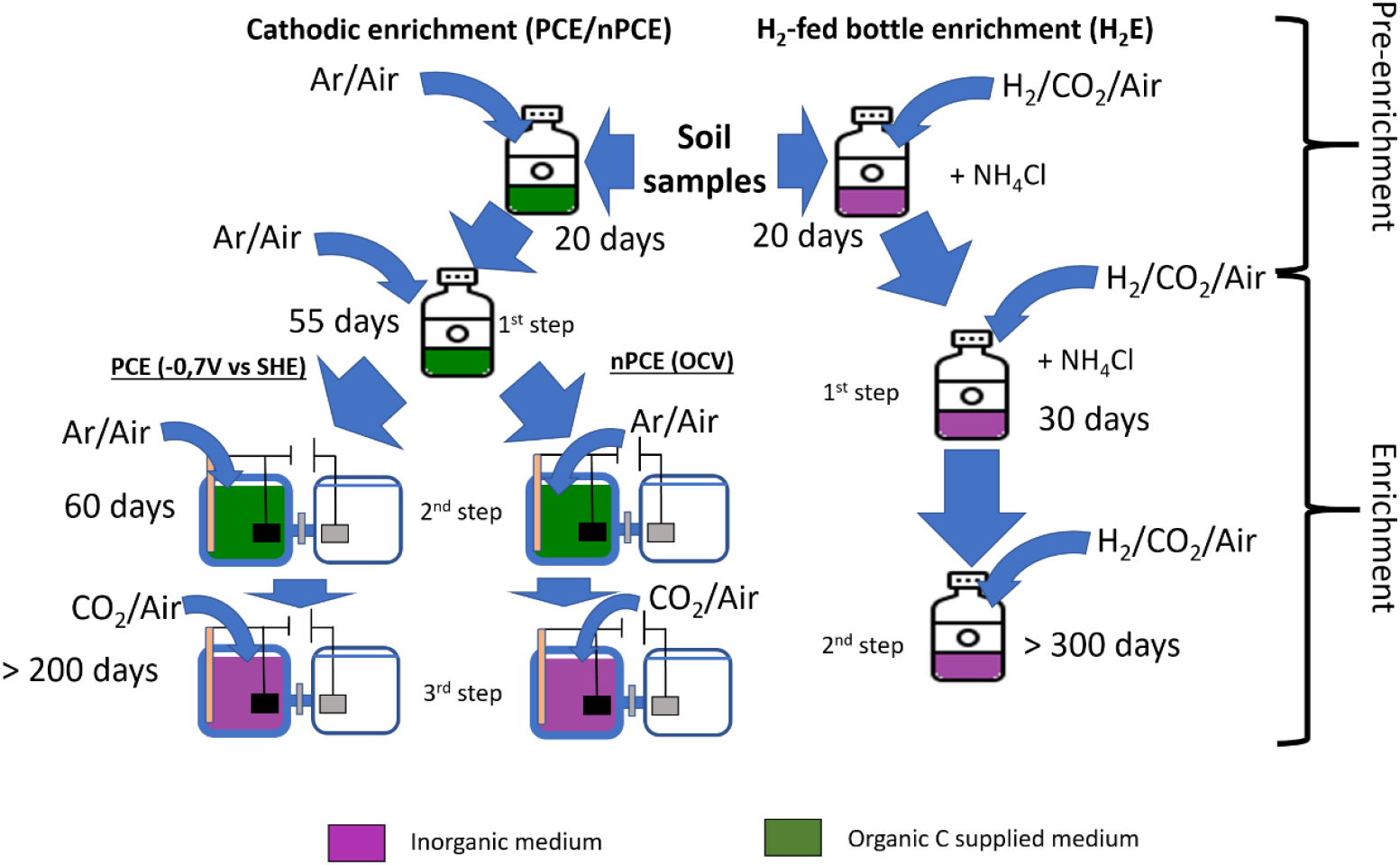
Diagram of the enrichment process

In <u>the second enrichment method, the H_2_-fed enrichment (H_2_E)</u>, autotrophic bacteria were enriched in inorganic medium supplemented with H_2_ as sole electron source. These enrichments on H_2_ were obtained by pre-enriching in strict autotrophic bacteria using 50 ml of inorganic medium in a 120 ml bottle. The headspace consisted of a mixture of 75/15/8/2 (v:v:v:v) H_2_/CO_2_/N_2_/O_2_ at an initial pressure of 1.5 bar (absolute). Two two-weeks batches (30 days) were performed with NH_4_Cl (1g/L) as nitrogen source in the first stage of this enrichment as seen in Figure 1. This nitrogen source was then replaced by N_2_ as only N source to enrich N_2_-fixing bacteria in the second stage of enrichment. Centrifugation (10min, 7500 RPM, ∼7500g) of 80% of the initial medium (40mL) was done at each subculture every 15-20 days. The pellets obtained after centrifugation were suspended in 5mL of sterile medium before being used for subculturing. Table 1 presents a summary of the different procedures used.

**Table 1.**
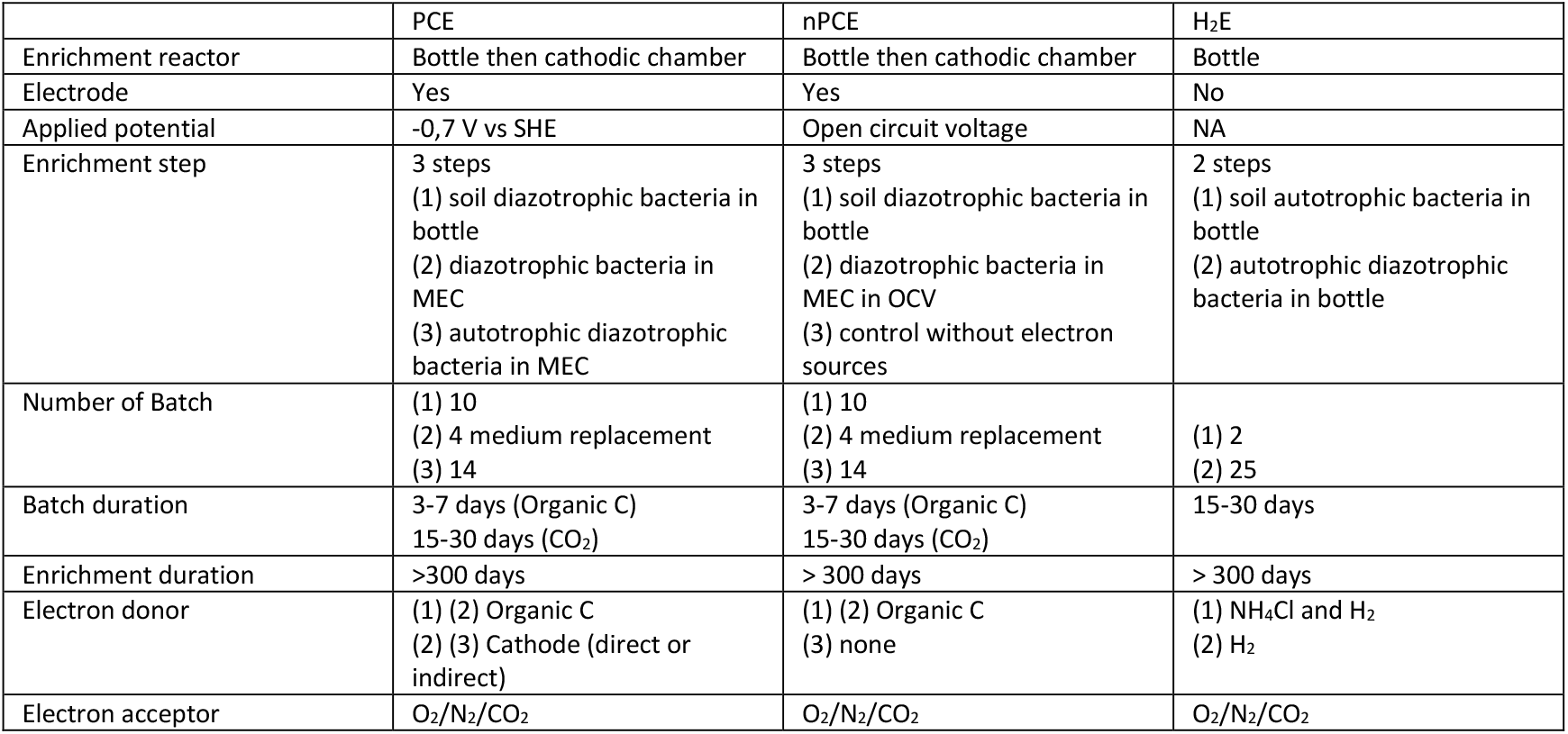
Comparison between each enrichments procedure used in this work.

### Medium analyses

NH_4_^+^, NO_2_^-^ and NO_3_^-^ concentrations were measured using a Gallery+ sequential analyzer (Thermo Fisher Scientific). Two mL of samples were centrifuged at 13500 RPM (∼12300g) and then filtered to 0.2μm with nylon membranes before being stored at 4°C. The remaining pellets were stored at −20°C and used for community analysis. VFAs and other carbon compounds were measured on Clarus 580 GC equipped with FID and a Dionex UltiMate 3000 HPLC as described elsewhere (Carmona-Martínez et al., 2015; Moscoviz et al., 2019)

Total nitrogen was measured using a CHNS Flashsmart elemental analyzer (Thermo Fisher Scientific). The sample (approximatively 2.5 mg) was weighed and was introduced into the oxidation/reduction chamber of the analyzer. 200 mL of medium were sampled at each medium change. These samples were dried for 4-5 days at 60°C. The samples were then freeze-dried and then grounded with a mortar. Two to four mg of each sample was used in the CHNS analyzer. The nitrogen content was then compared to the dry weight measured before freeze-drying to determine the mass of nitrogen in the medium. No CHNS analysis was performed on H_2_ –based enrichment due to a low culture volume (50mL).

Nitrogen present in the biomass (biofilm and planktonic) was estimated from quantification of 16S rDNA with qPCR. The rrnDB-5.7 database was used to estimate the actual bacterial amount from 16S rDNA qPCR using theoretical 16S rDNA copies/genome per strain, genus or family given by the database and sequencing results from our communities (Stoddard et al., 2015). Then, the theoretical calculated number of bacteria was used to determine the nitrogen content in the biomass using the theoretical average dry mass of an *Escherichia coli* cell of 216×10^−15^ g/bacterium and with a theoretical relative mass of nitrogen in microbial biomass, ie. 11.4% according to the average biomass formula CH_1.8_O_0.5_N_0.2_ (Heldal et al., 1985; Loferer-Krößbacher et al., 1998). The nitrogen present in the biomass was therefore estimated using the formula below:

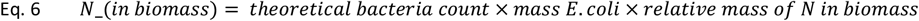

With *N*_*in biomass*_ the nitrogen concentration in the medium from biomass in mg_N_/L, the *theoretical bacteria count* based on the calcul of concentration in bacteria with results of qPCR of 16S rDNAfor each enrichment (number of bacteria/L), *mass E. coli* a constant of 2.16 10^−10^ mg/cell_*E. coli*_ and *relative mass of N in bacteria* is 11.4% of the dry mass.

### Gas analysis and acetylene reduction assay (ARA)

CO_2_, H_2_ and N_2_ used in the headspace of the enrichments were provided at laboratory grade. Pure ethylene was also supplied for calibration of the gas chromatography ethylene measurement.

Headspace compositions and pressures were analyzed every 1-2 days. The pressure was manually measured with a Keller LEO 2 manometer (KELLER AG, Wintherthur, Switzerland). Gas analyses were carried out on a Perkin Elmer Clarus 580 GC equipped with RT-Q-Bond and RT-Msieve 5Å columns with a TCD allowing the quantification of H_2_, CO_2_, N_2_, O_2_ and CH_4_ with Ar as carrier gas as described by A. Carmona-Martínez (Carmona-Martínez et al., 2015). Acetylene and ethylene were measured using a Perkin Elmer Clarus 480 GC equipped with RT-U-Bond and RT-Msieve 5Å columns with TCD and He as carrier gas as described in a previous work (Carmona-Martínez et al., 2015).

The acetylene reduction assay (ARA) was performed to quantify the rate of N_2_ fixation using the ability of nitrogenases to reduce acetylene to ethylene. This reaction occurs at a rate proportional to the rate of N_2_ fixation according to the theoretical ratio C_2_H_2_:N_2_ (3:1) (Bergersen, 1970). The ARA was performed only after 18 batch cycles for H_2_-fed enrichment (H_2_E) and 11 batch cycles in polarized cathode enrichment (PCE) fed with CO_2_ (340 days). Acetylene concentration were calculated as follow :

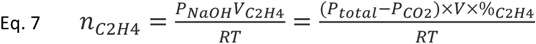

With (*P*_*total*_ the pressure in headspace, *P*_*CO*2_ the partial pressure of CO_2_ measured without CO_2_ trap, *V* the volume of gas in headspace, %_*C*2*H*4_ the part of C_2_H_4_ measured by GC-TCD in headspace, R the perfect gas constant and T, the temperature of the reactor.

And Acetylene production rate were calculated following next equation:

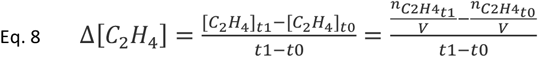

With Δ[*C*_2_*H*_4_] the rate of C_2_H_4_ production in μmol/L.d, *t0* the last measure without C_2_H_4_ observed, *t1* the first measure of C_2_H_4_ in headspace and V the volume of liquid which is constant between two ARA measurement.

A specific N_2_-fixing activity was then calculated with the N_2_-fixing bacteria measured by quantification of the *nifH* gene used as a marker for these bacteria. This specific activity corresponds to the rate of C_2_H_4_ produced per bacteria capable of fix N_2_ measured by qPCR of *nifH* gene. The acetylene used for the acetylene reduction assay (ARA) was obtained by adding calcium carbide (CaC_2_) in water and recovered in a bag. The acetylene concentration in the bag was then measured by gas chromatography. Gas from the bag was added to each enrichment to reach a composition of 10% V/V acetylene and an equivalent amount of gas was removed from the headspaces. Ethylene production was then daily monitored for 7 days in the microbial electrolysis systems (PCE and nPCE) and 15 in the H_2_E bottles by the Perkin Elmer Clarus 480 GC with TCD. To ensure separation of ethylene from CO_2_ on the RT-U-Bond column, a CO_2_ trap with sodium hydroxide (6M NaOH) was used at the time of sampling and is taken into account in the calculations. After the ARA method was completed, the headspaces were flushed with N_2_ and the gas recycling system was changed.

### Community sequencing and biomass quantification

The microbial communities were quantified using the 16S rDNA qPCR to determine the total bacterial concentration and *nifH* gene qPCR for N_2_-fixing bacteria. In parallel, 16S rDNA sequencing was performed to identify major members of each community. This sequencing was also necessary to convert the amount of 16S rDNA to total bacteria using the rrnDB-5.7 database. To analyze the communities present in suspension, 1.8 mL of sample were collected for qPCR. For the cathodes, 1 cm^2^ was recovered at several times. The piece of carbon felt was then chopped with a sterile scalpel before being immersed in 20 mL of sterile inorganic media. 1.8 mL was then recovered after shaking 20 mL of medium to resuspend as much biomass as possible. These samples were then centrifuged 10 min 13500 RPM (12340g). The supernatant was discarded and the pellets retained for DNA extractions. After qPCR, the concentrations measured in the electrode samples were expressed considering the volume of medium.

Genomic DNA was extracted using the PowerSoil™ DNA Isolation Sampling Kit (MoBio Laboratories, Inc., Carlsbad, CA, USA) according to the manufacturer’s instructions. The qPCR amplification program was performed in a BioRad CFX96 Real-Time Systems C1000 Touch thermal cycler (Bio-Rad Laboratories, USA). For the analysis of total bacteria, primers 330F (ACGGTCCAGACTCCTACGGG) and 500R (TTACCGCGGCTGCTGGCAC) were used. For the bacteria qPCR mix: SsoAdvanced™ Universal SYBR Green Supermix (Bio-rad Laboratories, USA), primer 330F (200 nM), primer 500R (200 nM), 2 μL of DNA, and water were used to a volume of 12 μL. The qPCR cycle was as follows: incubation for 2 min at 95°C and 40 cycles of dissociation (95°C, 10 s) and elongation (61°C, 20 s) steps. The results were then compared to a standard curve to obtain the copy number of the target in the sample. Both the 16S rDNA concentration of the PCE media and the cathodes are considered in the calculation of the total 16S rDNA concentration of the PCE. These concentrations are used as an indicator of the biomass present and the use of a database of the number of 16S operons per bacterial genome was used to estimate the actual amount of bacteria.

The presence of N_2_-fixing bacteria was monitored by qPCR of the *nifH* gene of the Fe-Fe subunit of nitrogenases (Dos Santos et al., 2012; Gaby & Buckley, 2012). The *nifH* gene is known as marker of N_2_ fixing bacteria, common to all nitrogenases and is used for their quantification because of its necessary presence for the fixation of N_2_ (Dos Santos et al., 2012; Gaby & Buckley, 2012). All qPCR amplification programs were performed in a BioRad CFX96 Real-Time Systems C1000 Touch thermal cycler (Bio-Rad Laboratories, USA). The primers PolF-TGCGAYCCSAARGCBGACTC and PolRmodify reverse-AGSGCCATCATYTCRCCGGA were used (Poly et al., 2001). The mixture: 6μl SsoAdvanced™ Universal SYBR Green Supermix (Bio-rad Laboratories, USA), F primer (500 nM), R primer (500 nM), 2 μL of DNA and water was used up to a volume of 12 μL. The qPCR cycle was as follows: incubation for 2 min at 95°C and 40 cycles of dissociation (95°C, 30 s) and elongation (60°C, 30 s) steps. Then, the results were compared to a standard curve to obtain the number of copies of the target in the sample. These two quantifications allow us to calculate the ratios of N_2_-fixing bacteria per total bacteria of the enrichments at different points to track the enrichment of N_2_-fixing bacteria. This ratio can also help us to derive hypotheses on the functioning of our communities that can be completed by the analysis of the communities during sequencing.

After quantification, our enriched communities were sequenced according to their 16S rDNA and the results are available on NCBI repository PRJNA976100, Biosample SAMN28447998-SAMN28448066. The V3-V4 region of the 16S rDNA was amplified using universal primers as reported elsewhere (Carmona-Martínez et al., 2015). The PCR mixture consisted of MTP Taq DNA Polymerase (Sigma-Aldrich, Germany) (0.05 u/μL) with enzyme buffer, forward and reverse primers (0.5 mM), dNTPs (0.2 mM), sample DNA (5-10 ng/μL), and water to a final volume of 60 μL. 30 cycles of denaturation (95°C, 1 min), annealing (65°C, 1 min), and elongation (72°C, 1 min) were performed in a Mastercycler thermal cycler (Eppendorf, Germany). A final extension step was added for 10 min at 72 °C at the end of the 30th amplification cycle. PCR amplifications were verified by the 2100 Bioanalyzer (Agilent, USA). The GenoToul platform (Toulouse, France http://www.genotoul.fr) used an Illumina Miseq sequencer (2 × 340 bp pair-end run) for the sequencing reaction. The raw sequences obtained were analyzed using bioinfomatic tools. Mothur version 1.39.5 was used for cleaning, assembly and quality control of the reads. Alignment was performed with SILVA version 128 (the latter was also used as a taxonomic contour).

Communities sequenced on pre-enrichment bottles were used as initial community of the enrichment cultures. For PCE and nPCE, sequenced communities came from the biofilm formed on electrodes. Two replicates per potential were used. For H_2_E bottles, 6 bottles were used for sequence analysis and qPCR. For pre-enrichment, sequences corresponded to each soil samples (leguminous, forest, compost) and a mix of them.

### Data analysis

All results were analyzed using R (4.2.0) and Rstudio (2022.07.1) for calculations and graphics. The Tidyverse package was used for data manipulation(*Tidyverse*, n.d.). The packages ggplot2, ggpubr, scales, cowplot, corrplot and palettetown were used for the graphical representations. Visual representation of bacterial relative abundances was performed with the phyloseq package (McMurdie, 2011/2023). Inkscape software was also used to edit the graphs when necessary. The uncertainties shown for the values presented are standard deviations. All data and scripts used here are available online (Rous, 2023).

## Results and discussion

### Nitrogen fixation after 340 days of enrichment

N_2_ fixation was quantified at the day 340 of the enrichment using acetylene reduction assays (ARA). This assay was performed in H_2_-fed bottle enrichments (named ‘H_2_E’), in polarized cathode enrichment (named ‘PCE’) and in the non-polarized cathode enrichments as controls (named ‘nPCE’). As shown in Figure 2, the ARA results confirmed the N_2_ fixation capacity of the enriched communities (Bergersen, 1970). This indicates that the cathode and/or H_2_ was used as electron sources for N_2_ fixation both in PCE and in H_2_E bottles. The average rates were similar in both enrichment methods with 32±17 μmolC_2_H_4_/L.d in PCE and 36±13 μmolC_2_H_4_/L.d in H_2_E bottles. The PCE corresponding N_2_ fixation rates ranged from 0.12 mg_Nfixed_/L.d (minimum) to 0.51 mg_Nfixed_/L.d (maximum), which is consistent with the rate of 0.2 mg_Nfixed_/L.d estimated by Rago et al. (2019) and also with N_2_ fixation rates reported for soil bacteria (Hardy et al., 1973; Kifle & Laing, 2016; Rago et al., 2019). Despite these significant N_2_ fixation rates and the long duration of the experiments, the rate of ammonium production in solution remained lower than 0.07 mg_N_/L.d at the day 340 (Table 2), indicating that most of the fixed N_2_ was probably rapidly used by bacteria.

**Table 2.** The average ammonium production rates, the number of *nifH* gene copies, the number of 16S gene copies, and the *nifH*/16S ratio were assessed after 340 days for the three experimental configurations. Average values were measured on the last batch of 21 days, before 340 days of enrichment for the two polarized cathode enrichment (PCE), the two non-polarized Cathode Enrichments (nPCE) and the six H_2_ enrichment bottles (H_2_E)

|  | PCE | nPCE | H<sub>2</sub>E |
| --- | --- | --- | --- |
| N-NH<sub>4</sub><sup>+</sup> mg/L.d | 0.07±0.01 | 0.07±0.01 | 0.04±0.03 |
| *nifH* gene copies copies/mL | 7.8±9.5 10<sup>7</sup> | 2.5±2.8 10<sup>6</sup> | 8.3±2.2 10<sup>5</sup> |
| 16S rDNA gene copies copies/mL | 1.7±1.9 10<sup>9</sup> | 1.4±0.2 10<sup>8</sup> | 4.6±1.5 10<sup>6</sup> |
| *nifH*/16S | 3.8 % | 1.7 % | 19.0 % |

**Figure 2.**
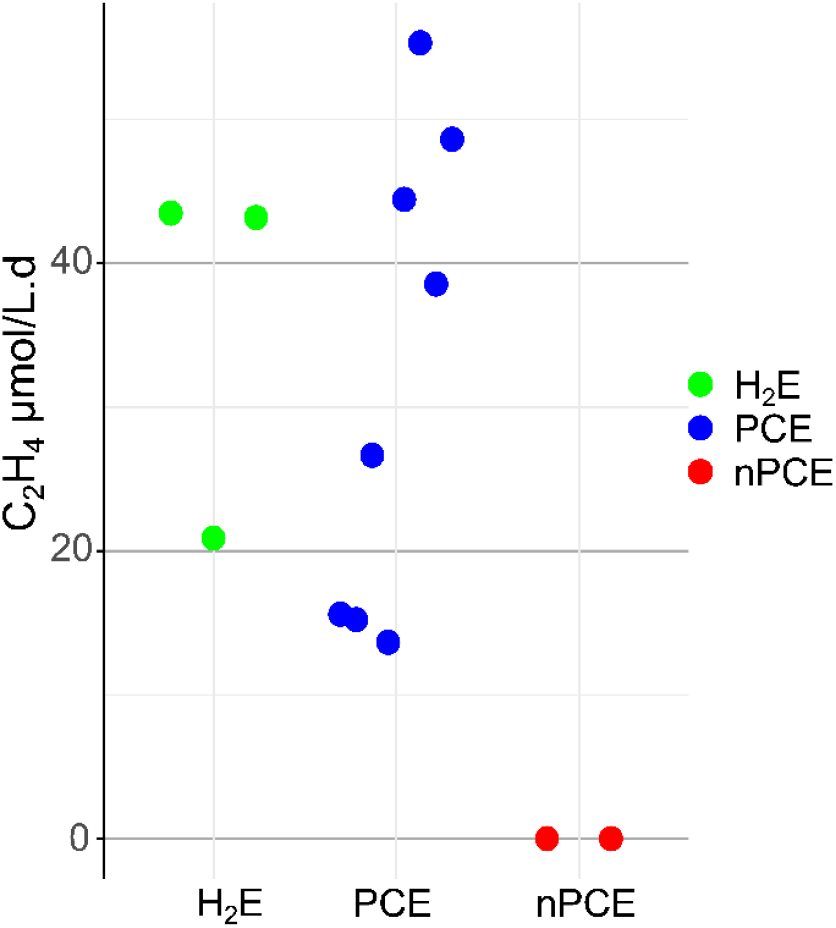
Reduction rate of acetylene in μmol C_2_H_4_/L.d in the different reactors after 340 days of operation. or H_2_ condition, 3 bottles were used for the acetylene reduction assay with one injection for each bottle, i.e. 3 measurements. For PCE, 2 reactors were used and 4 injections were made to validate the repeatability of the measurement when C_2_H_4_ which gives 8 measurements for each of these two conditions. For nPCE, 2 reactors were used which gives 2 measurements for each conditions.

The ability of the microbial communities to fix N_2_ was also assessed by qPCR of the *nifH* gene (Dos Santos et al., 2012; Pogoreutz et al., 2017). The amounts of N_2_-fixing bacteria after 340 days of enrichment are reported in Table 2. The average *nifH* gene concentration in PCE was estimated at 7.8 10^7^ copies_*nifH*_/mL, two orders of magnitude higher than the average concentration of 8.3 10^5^ copies_*nifH*_/mL measured in the H_2_E enrichment bottles. This observation was surprising since N_2_-fixation rates were similar in both configurations (Figure 2). This suggests that the fixation rate per *nifH* copy was much higher in H_2_E than in PCE. The estimated specific activities per *nifH* copy were indeed of 0.2±0.3 μmol_C2H4_/10^8^ copies_*nifH*_.d in the PCE and 2.1±0.7 μmol_C2H4_/10^8^ copies_*nifH*_.d in H_2_E bottles (Table 2). Furthermore, the comparison between *nifH* gene and 16S rDNA copy numbers gives an idea of the proportions of N_2_ fixing bacteria in each microbial community. Interestingly, this proportion was 18% in H_2_E bottles that was four times higher than the value of 4.6% in PCE (Table 2). Therefore, N_2_-fixing bacteria constituted a smaller proportion of the bacterial populations in PCE than in H_2_E and only a small proportion of the bacteria participated to N_2_ fixation in the PCE. In nPCE controls, the biomass was higher than in H_2_E and the *nifH*/16S ratio lower (Table 2). This higher biomass concentration was likely due to the enrichment period in presence of organic C (day 0 to 115 including 60 days in BES) for the PCE and nPCE enrichments. NH_4_^+^ in nPCE was also observed at a rate of 0.07 mg/L.d as presented in Table 2 but without acetylene accumulation, meaning that no N_2_ fixation occurred. In absence of electron source, nPCE enrichment communities could have been maintained through cryptic growth. The presence of NH_4_^+^ in the nPCE was likely related to cell lysis since no measurable fixation was detected by ARA even though significant biomass production was observed.

### Current density and Autotrophic enrichment in polarized cathode enrichment

The average current density for the two PCE over experimental time is shown in Figure 3. As the current measured at the cathode was negative by convention, a more negative corresponded to a higher reduction activity. Regarding the current densities in the abiotic systems, the average current densities were measured at −0.75 A/m^2^ for two times four days with an organic C source and −1.1 A/m^2^ for 16 days with only CO_2_ as the carbon source.

**Figure 3.**
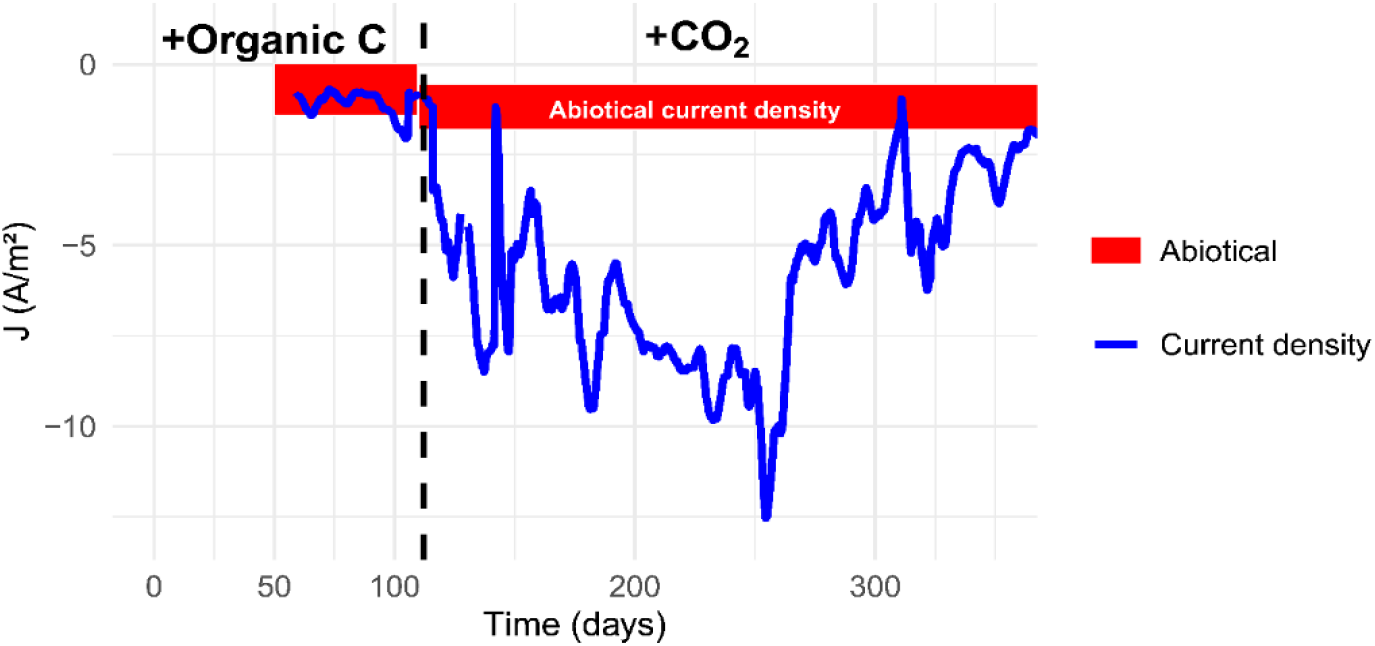
Mean current density measured for the two PCE (blue line). Levels shown in red correspond to theoretical mean current density and standard deviation estimated from two abiotic electrochemical systems current densities. For abiotic electrochemical system, one batch of 2 days were made with Organic C and one batch of 16 days with CO_2_. The peaks observed are due to the batch operation of the PCE with disturbances each time the medium was renewed. Power failures occurred at 230 days and 260 days. The time indicated on the x axis corresponds to the experimental time starting at day 0 of the enrichment where soil samples were first introduced in the bottles with a medium containing organic C. Day 55 corresponds to the start of microbial electrochemical system with the precultured communities. The dashed line on day 110 corresponds to the passage on CO_2_ as the sole carbon source in PCE. A 5-day curve smoothing was applied on PCE current denstity curve.

The average current density measured in the PCE was not different from the current density measured in the abiotic control (approximately −1 A/m^2^ with organic C supply) for the first 45 days of operation (days 55 to 100 of the enrichment). An increase up to −2 A/m^2^ appeared between 100 and 115 days of enrichment in PCE. CO_2_ was then used as sole carbon source after this increase appeared. Following this change in carbon source, a sharp increase in current density to −5 A/m^2^ was observed in the PCE with regards to the range of current densities in the abiotic controls (approximately −2 A/m^2^ on CO_2_)(Figure 3). The higher current density was assumed to be associated to the electroactive activity of electron uptake by enriched bacteria. The current density in the PCE then continuously increased until day 250 of the enrichment to reach a value of −15 A/m^2^.

The high current density observed after 250 days of enrichment indicated a high redox activity linked either to hydrogen evolution, oxygen reduction, or possibly direct electron transfer. As proposed by Z. Zaybak et al. (2013), the high activity was probably resulting from a high metabolic activity in the biofilm with significant microbial catabolic process (Zaybak et al., 2013). Compared to the current densities obtained Rago et al. (2019) in the order of magnitude of −10 mA/m^2^ at the same potential (−0.7 vs SHE), the current densities observed here (−5 to −10 A/m^2^) were about 1000 times higher. These current density levels are close to those measured by Zhang et al. (2022) who reported a maximum of −10 A/m^2^ at the same applied potential (Zhang et al., 2022).

After 230 days, power failures occurred, interrupting temporarily the current supply to the cathodes. An important decrease of the current density was observed afterwards, down to −5 A/m^2^ after 260 days and −3 A/m^2^ after 320 days. The lower current density reflected a change in the functioning of the microbial communities, leading to less electron exchange with the cathode.

### Biomass quantification

Bacterial biomass was monitored in the enrichments by measuring 16S rDNA concentrations by qPCR (Figure 4). At day 131 (18 days after switching from organic C to CO_2_), the average 16S rDNA concentrations measured in the polarized cathode enrichment (PCE) and in the non-polarized cathode enrichment (nPCE) controls were 4.6±0.4 10^9^ and 4.2±1.2 10^9^ copies16SrDNA/mL, respectively. These concentrations corresponded to 9.4±1.0 10^8^ and 8.0±1.5 10^8^ bacteria/mL, respectively, as presented in Figure 4. These bacteria concentrations resulted from the first enrichment phase with organic C. During this phase, organic substrates were used as carbon and electron sources for biomass growth in both configurations (PCE and nPCE). At day 214, the 16S rDNA concentration in the PCE was used to calculate a concentration of 3.3±2.1 10^9^ bacteria/mL, corresponding to a biomass increase by a factor of 3.5 between 131 and 214 days (Figure 4). At the same time, the bacterial concentration dropped in nPCE from 8.0±1.5 10^8^ bacteria/mL to 3.0±0.8 10^8^ bacteria/mL. This drop was explained by the lack of available energy source for growth, which led to a sharp decrease of the bacterial populations. At day 214, the 16S rDNA concentration in PCE was therefore 11-fold higher than in nPCE controls. This difference is consistent with the difference reported by Rago *et al*. between polarized and non-polarized conditions, with electroactive biocathodes enriched in autotrophic diazotrophic bacteria (Rago et al., 2019). These results suggested that the enriched microbial communities were able to use the electrodes polarized at −0.7 V vs. SHE as sole electron sources to grow while fixing N_2_.

**Figure 4.**
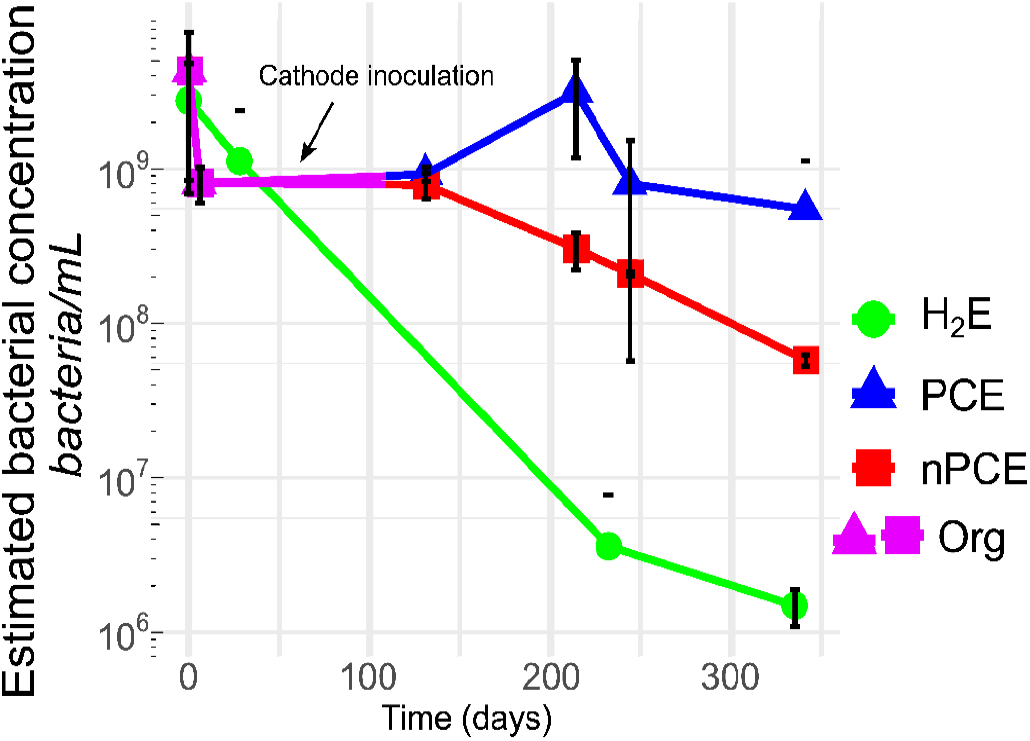
Bacteria concentrations over time in the different enrichments calculated from 16S rDNA qPCR quantifications in bulk and biofilm. Green disks correspond to H_2_ enrichments in bottles, blue triangles correspond to polarized cathode enrichment (PCE), red squares correspond to controls in non-polarized cathode enrichment (nPCE). The partially purple symbols marked Org correspond to the first phases of enrichment with organic C for the PCE and nPCE. The arrow indicates the transition from bottle enrichments to cathode enrichments in microbial electrochemical systems for PCE and nPCE. Error bars correspond to the calculated standard deviation.

In the H_2_-fed enrichment (H_2_E) bottles, the 16S rDNA concentration steadily decreased along the experiment. The concentration decreased from 1.1±1.3 10^9^ bacteria/ml at the beginning of the enrichment down to 1.3±0.3 10^6^ bacteria/ml after 340 days (Figure 4). These concentrations appear lower than the biomass concentrations observed in the PCE medium at the same time of enrichment, ie. 4.5±5.9 10^6^ bacteria/mL at day 244 and 8.6±8.8 10^6^ bacteria/mL at day 340. These results confirm that bacterial growth was higher on the cathodes than in an H_2_ supplied environment. It was therefore concluded that the PCE provided more favorable environment for biomass growth than H_2_ fed bottles as the surface provided by the electrode was likely favorable for biofilm growth.

We also calculated the *nifH*/16S ratio representing the part of bacteria able to fix N_2_ among the total bacteria. A ratio of 0.0006 of *nifH* gene copies per 16S rDNA copy was measured for samples at the very beginning of enrichment, both for H_2_E and PCE. After 131 days, corresponding to the switch to CO_2_ as sole C-source, this level increased to 0.03 in PCE and 0.02 in nPCE control. These results are consistent with an enrichment in diazotrophic bacteria during the enrichment phase on organic carbon (Bowers et al., 2008). The bacterial enrichment in nPCE was likely possible as the organic C was used by the bacteria as energy source. After 214 days, the level decreased to 0.02 in PCE but remained higher than the level at the beginning of enrichment. This variation suggests interactions within the community that favored the growth of non-N_2_ fixing bacteria after the shift to CO_2_ as sole C-source. After 340 days, the ratio of *nifH* to 16S rDNA was 0.04as presented in Table 2. In parallel, a ratio of *nifH* to 16S rDNA of 0.90 was measured for H_2_E at 244 days. Therefore, most of the bacteria were able to fix N_2_ in H_2_E bottles, confirming the efficient enrichment in diazotrophic bacteria (Bowers et al., 2008). Given the loss of biomass observed in H_2_E during the experiment (Figure 4), this high ratio corresponded likely to the surviving bacteria that were selected on their ability to fix N_2_. The ratio measured in these H_2_E then decreased down to 0.19, suggesting a decrease in N_2_-fixing bacteria in biomass.

As previously mentioned, after 230 days, power failures occurred and interrupted the polarization of the electrodes. These interruptions impacted the microbial communities with a decrease in biomass concentration to 8.1±7.6 10^8^ bacteria/mL at 244 days and 5.5±6.0 10^8^ bacteria/mL after 340 days compared to the concentration of 3.3 10^9^ bacteria/mL measured at 214 days. At the same time, the *nifH*/16S rDNA ratio increased up to 5%, indicating that N_2_-fixing bacteria were more resistant. Nevertheless, a decrease was observed in *nifH* quantities, from 2.3 10^8^ copies_*nifH*_/mL after 214 days to 7.8 10^7^ copies_*nifH*_/mL after 340 days.

### N quantification and coulombic efficiency

Total N contents of the different experiments are shown in Figure 5a. The total N corresponded to the sum of the nitrogen measured in the liquid phase by N-ion concentration analysis (N-NH_4_^+^, N-NO_3_^-^, N-NO_2_^-^), in the medium by CHNS elemental analysis for PCE and nPCE, in the suspended biomass from qPCR results only for H_2_E where the dry mass was not measured, and on the electrode based on the bacterial concentrations. The total N concentration was estimated after 131 days of enrichment (ie. before the shift to CO_2_ as sole C source) at 25.5±0.4 mg_N_/L and 25.8±4.7 mg_N_/L in the PCE and in the nPCE controls, respectively. After 214 days of enrichment, total N increased up to 87.6±56.1 mg_N_/L in the PCE with regards to the low value of 14.8±7.0 mg_N_/L in the nPCE controls. Ammonium represented only a small fraction of the total N in the PCE. The maximum ammonium concentration observed at a batch end in the PCE was 1.5±0.3 mg_N_/L at the day 340 of the enrichment in comparison with the maximum value of 4.5±0.4 mg_N_/L found in nPCE control after 131 days (1.4 mg_N_/L at the day 340)(Figure 5b). The average N fraction in the form of ammonium was therefore of 12% in nPCE and only 1.6% in PCE. It was thus assumed that the higher level of ammonium in nPCE was related to the decay of biomass in absence of electron sources. In counterpart, the ammonium produced by N_2_ fixation in the PCE was likely directly used for protein synthesis as suggested elsewhere (Bueno Batista & Dixon, 2019; Temple et al., 1998). In addition, nPCE N-NO_3_^-^ concentrations were 5 to 15 times lower than N-NH_4_^+^ concentrations with maximum values of 0.9 mg_N_/L N-NO_3_^-^ in the nPCE control and 0.2 mg_N_/L N-NO_3_^-^ in PCE. An average concentration of 0.1±0.2 mg_N_/L N-NO_3_^-^ in H_2_ E bottles was also observed. NO_2_^-^ concentrations were negligible.

**Figure 5.**
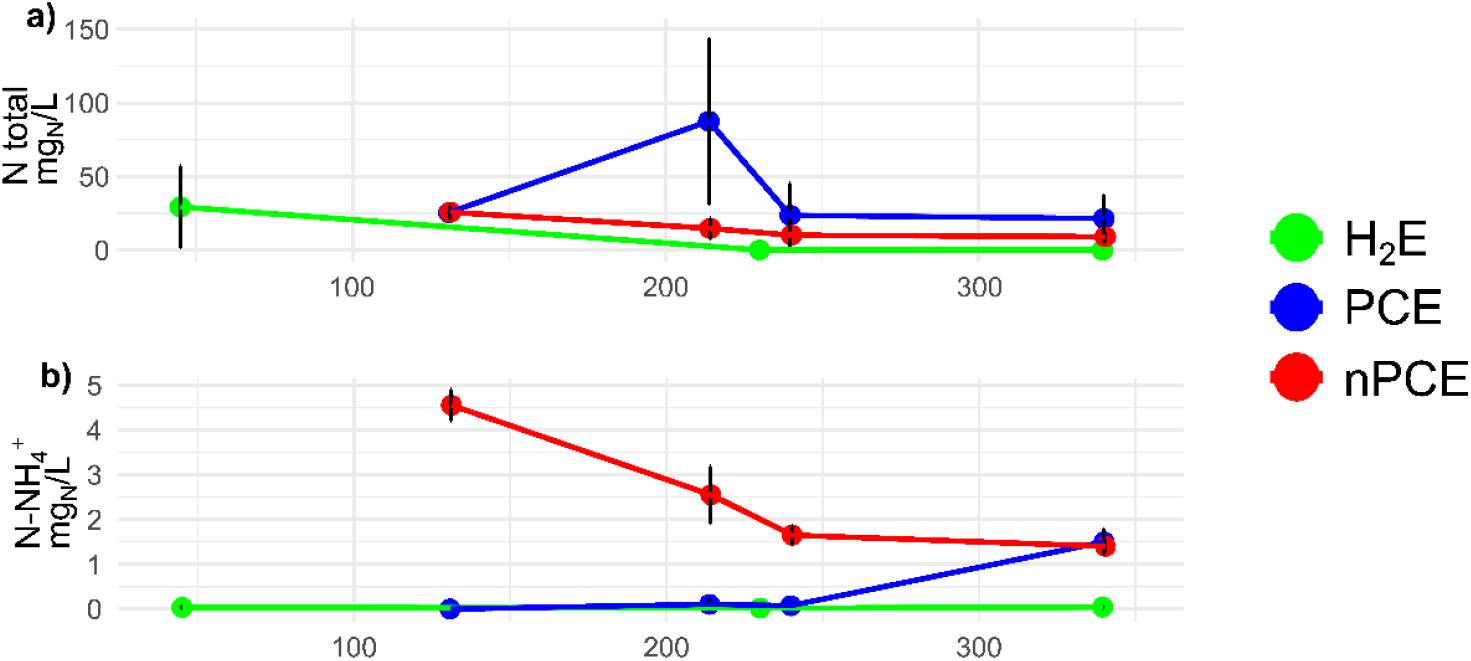
(a) Total N concentration based on the sum of (1) N estimated from biomass measurement (suspended biomass and biofilm), (2) N content in ionic forms (N-NH_4_^+^, N-NO_3_, N-NO_2_) and (3) N measured in the dry weight of the medium of the polarized cathode enrichment (PCE) and non-polarized cathode enrichment (nPCE) and (b) N-NH_4_^+^ concentration in H_2_ enrichment (H_2_E), PCE and nPCE.

Using our method of N mass estimation on biomass lost as presented in Figure 4 for nPCE, the total concentration of N lost by biomass would be estimated around 130 to 200 mgN/L depending on the community (2 nPCE). This loss would then be equivalent to rates of 0.6 to 0.9 mgN/L.d released by this the biomass on average along the enrichment. Assuming a loss of a constant portion of biomass, a release rate of 0.1 mgN/L.d was estimated as average for the batch ending at 340 days, close to the 0.07 mg/L.d presented in Table 2. This result supported our hypothesis that NH_4_^+^ release was linked to biomass loss in nPCE enrichments after organic C addition was stopped.

The H_2_ enrichment bottles (H_2_E) showed an average N-NH_4_^+^ accumulation of 35.6±36 μg_N_/L after 131 days and an average of 0.1±0.2 mg_N_/L over the duration of enrichment. This concentration represented a small fraction of the total N at the very beginning of the enrichment, with an estimated concentration of 29.4±28 mg_N_/L. The N concentration decreased during the enrichment, which is consistent with the biomass loss as shown in Figure 4. H_2_E were therefore less efficient for N_2_ accumulation than polarized cathode enrichment, with a lower microbial biomass production.

Current densities and rates of acetate production, N_2_ fixation and biomass growth are shown in Table 3. Coulombic efficiencies associated with each reaction were estimated based on these results (Table 3). During the first 214 days of enrichment, 0.6 to 3.3% of the electrons were used for N_2_ fixation in the two PCE. In comparison, efficiencies of 0.5% and 20% for NH_4_^+^ synthesis was reported in two recent works carried out under similar conditions (Yadav et al., 2022; Zhang et al., 2022). As the amount of fixed N was highly dependent on biomass accumulation, negative results were obtained at day 244 when biomass started to decrease. The electrons used at the cathode for biomass synthesis during the first period (131 to 214 days) accounted for 2.8% and 17.3% in the two PCE. These high coulombic efficiencies were probably also associated to acetotrophic and acetogenic bacteria. Indeed, acetate produced using electron from the cathode could have been used by acetotrophic bacteria for growth, lowering coulombic efficiency of acetogenesis and increasing CE for bacterial growth. As acetogenic bacteria do not tolerate the presence of oxygen, dissolved O_2_ was very probably consumed in some part of the biofilm, leaving other parts in strict anaerobic conditions more favorable for acetogenic bacteria growth.

**Table 3.**
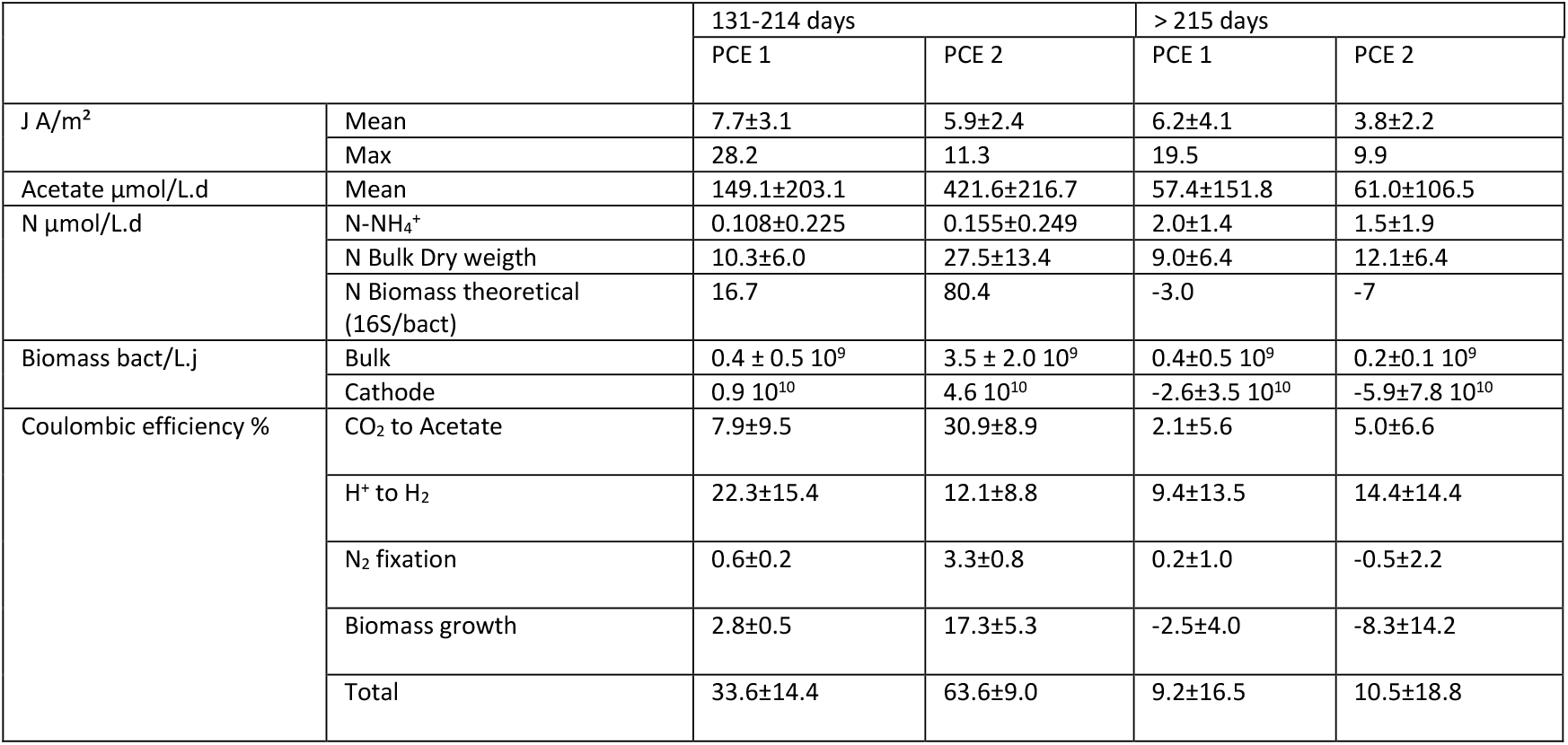
Current densities, rates and coulombic efficiencies for the two polarized cathode enrichment (PCE) over two different periods of current density with CO_2_ as sole carbon source. During the first period (131-214 days) current density increased, whereas during the second period (>215) current density decreased after several power outages (see Figure 3).

|  |  | 131-214 days |  | > 215 days |  |
| --- | --- | --- | --- | --- | --- |
|  |  | PCE 1 | PCE 2 | PCE 1 | PCE 2 |
| J A/m<sup>2</sup> | Mean | 7.7±3.1 | 5.9±2.4 | 6.2±4.1 | 3.8±2.2 |
|  | Max | 28.2 | 11.3 | 19.5 | 9.9 |
| Acetate µmol/L.d | Mean | 149.1±203.1 | 421.6±216.7 | 57.4±151.8 | 61.0±106.5 |
| N µmol/L.d | N-NH<sub>4</sub><sup>+</sup> | 0.108±0.225 | 0.155±0.249 | 2.0±1.4 | 1.5±1.9 |
|  | N Bulk Dry weight | 10.3±6.0 | 27.5±13.4 | 9.0±6.4 | 12.1±6.4 |
|  | N Biomass theoretical (16S/bact) | 16.7 | 80.4 | -3.0 | -7 |
| Biomass bact/L.j | Bulk | 0.4 ± 0.5 10<sup>9</sup> | 3.5 ± 2.0 10<sup>9</sup> | 0.4±0.5 10<sup>9</sup> | 0.2±0.1 10<sup>9</sup> |
|  | Cathode | 0.9 10<sup>10</sup> | 4.6 10<sup>10</sup> | -2.6±3.5 10<sup>10</sup> | -5.9±7.8 10<sup>10</sup> |
| Coulombic efficiency % | CO<sub>2</sub> to Acetate | 7.9±9.5 | 30.9±8.9 | 2.1±5.6 | 5.0±6.6 |
|  | H<sup>+</sup> to H<sub>2</sub> | 22.3±15.4 | 12.1±8.8 | 9.4±13.5 | 14.4±14.4 |
|  | N<sub>2</sub> fixation | 0.6±0.2 | 3.3±0.8 | 0.2±1.0 | -0.5±2.2 |
|  | Biomass growth | 2.8±0.5 | 17.3±5.3 | -2.5±4.0 | -8.3±14.2 |
|  | Total | 33.6±14.4 | 63.6±9.0 | 9.2±16.5 | 10.5±18.8 |

The H_2_ recovered in the headspace of the PCE accounted for 12 to 22% of the electrons supplied to the cathode as presented in Table 3. Therefore, H_2_ was not related to the biological activity and mostly resulted from an abiotic reaction at the cathode. In addition, in presence of O_2_ in the cathodic chamber, oxygen reduction reactions were expected with regards to the potential used in this study. Indeed, a two-electron reduction could have occurred, resulting in the production of hydrogen peroxide (H_2_O_2_), which can then undergo further reduction to form water (H_2_O) (Rozendal et al., 2009; Sim et al., 2015). Hydrogen peroxide was not measured, however, the amount of biomass found on the cathodes (Figure 4) suggested that the concentrations of hydrogen peroxide were sufficiently low to have minimal to no impact on the microbial community during the enrichment process. Nonetheless, a fraction of the electrons may have been lost through these oxygen reduction reactions, which could partially account for the low coulombic efficiencies observed in this study. Interestingly, a significant production of acetate was also observed. An average rate of 149.1 μmol/L.d and 421.6 μmol/L.d were measured in both PCE for the period from day 131 to day 214, as presented in Table 3. Acetate production almost stopped with the power failures with acetate measured only on one to two batches per PCE. This decrease correspond to rate of 61.0 μmol/L.d and 57.4 μmol/L.d of acetate in both PCE. Acetate production accounted for 7.9 and 39% of the cathodic electrons during the first period (up to 214 days) and for less than 5% after power failures. To explain the decrease in acetate production, biomass growth and power consumption, it was hypothesized that acetate production might have been due to a specific loss of members able to fix CO_2_, and more especially autotrophic bacteria using cathode as electron source (H_2_ or DET), within the enriched community. Thus, with acetate no longer being produced, heterotrophic bacteria did not have enough organic C to sustain their growth, causing their decrease. Therefore, autototrophic bacteria responsible for CO_2_ fixation and heterotrophic bacteria that could also be H_2_ dependent for N_2_ fixation were greatly affected, leading to a decrease in the reduction reactions at the cathode and subsequently in current density. In addition, acetate was not found in the H_2_E bottles, indicating the absence of acetogenesis and an important difference in microbial pathways and/or communities. As seen in the Table 3, electrons were retrieved in biomass production, N_2_ fixation products, H_2_ found in headspace and in CH_3_COOH product from CO_2_. These products were not sufficient to close the electron mass balance. The loss of electrons and the differences between cathodes of PCE 1 and PCE 2 was explained by side reactions, such as O_2_ reduction or biological reaction such as exopolysaccharide (EPS) production.

In comparison with the other works dealing with N_2_-fixing cathodic biofilms, Zhang et al (2022) showed a maximum of 40.5 mg/L of NH_4_^+^ in 4 days with mixed communities, in regard to a maximum of 0.8 mg/L NH_4_^+^ observed in Yadav et al (2022) and 6.31 mg/L NH _4_^+^ in 10 days for Li et al. (Li et al., 2022; Yadav et al., 2022; Zhang et al., 2022). When demonstrating the N_2_ fixation in MES, Rago et al (2019) showed a N_2_ fixation of 0.2 mgN/L.d in biomass and 5 10^9^ bacteria/L.d (Rago et al., 2019). In the present study, biomass production in biofilms was 2 to 10 times higher than in Rago et al. (2019), as was the nitrogen found in the biomass varied between 0.2 and 1 mg/L.d before 214 days (Rago et al., 2019).

### Microbial communities

16S rDNA sequencing was performed at the end of pre-enrichment, and at 214 or 232 days of enrichment in polarized cathode enrichment (PCE) and in H_2_-fed enrichment bottles (H_2_E), respectively. The sampling days were selected because they were associated to a high microbial activity (high current densities and high biomass concentrations). In H_2_E, the *nifH*/16S abundance ratio was also maximum (0.9) at day 232. Principal Component Analysis (PCA) was used to present the communities for each enrichment. Each reactors and bottles are presented as individuals and major families as variables in PCA presented in the Figure 6 and relative abundances are presented in the Figure 7.

**Figure 6.**
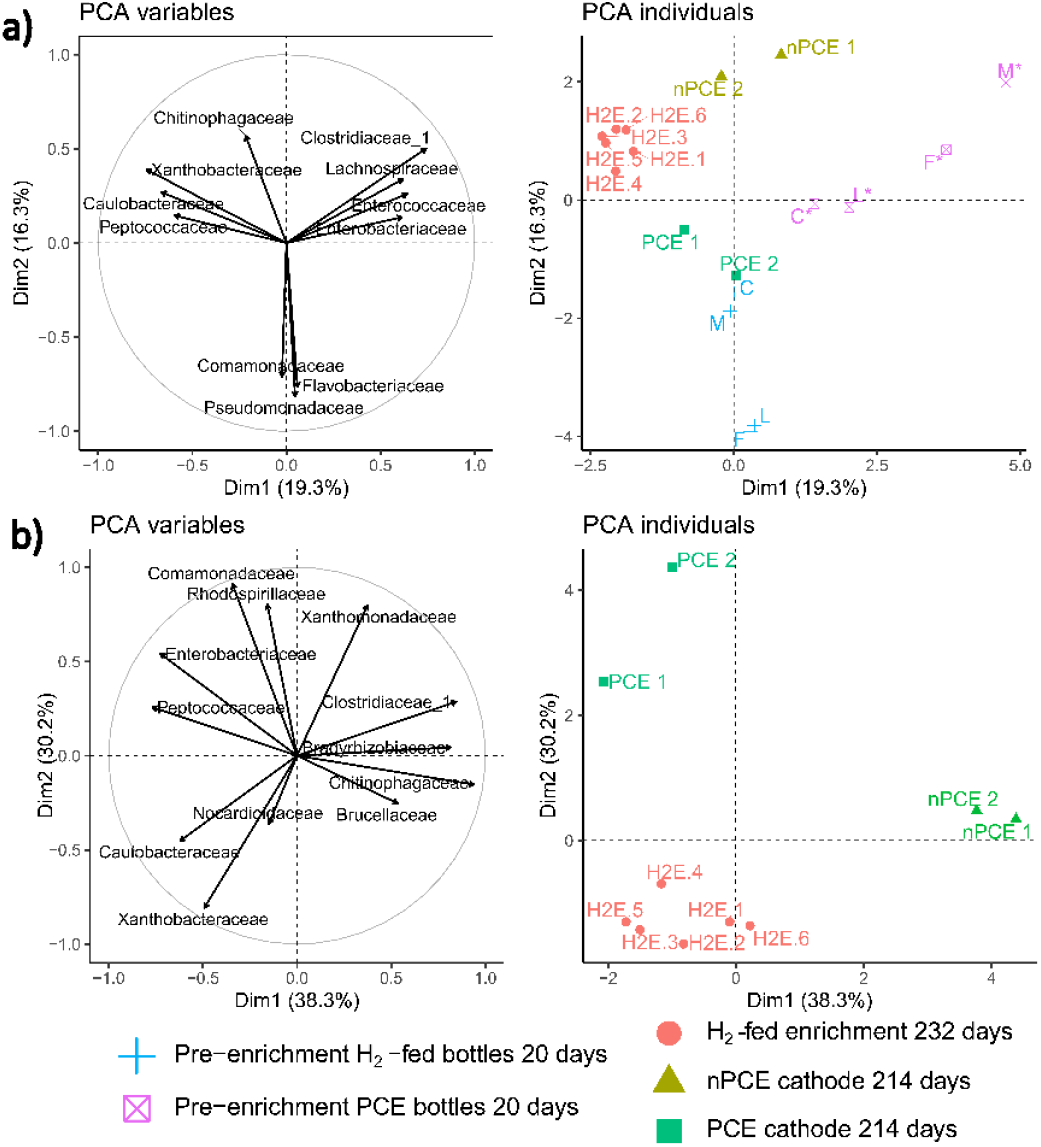
Results of the principal component analysis (PCA) performed on the microbial communities of a) pre-enrichments, H_2_-fed enrichment bottles (H_2_E) after 232 days and cathodes of PCE and nPCE enrichments after 214 days and b) H_2_-fed enrichment bottles (H_2_E) after 232 days and cathodes of PCE and nPCE enrichments after 214 days. Only families of the five major bacterial OTU in each sampled community were used for the analysis. The microbial communities in the pre-enrichment bottles are represented by the following abbreviations: F for forest soil, C for compost, L for the rhizosphere of leguminous plants, and M for a mix of all. Variables least close to the correlation circle are not displayed (cos2 < 0.2).

**Figure 7.**
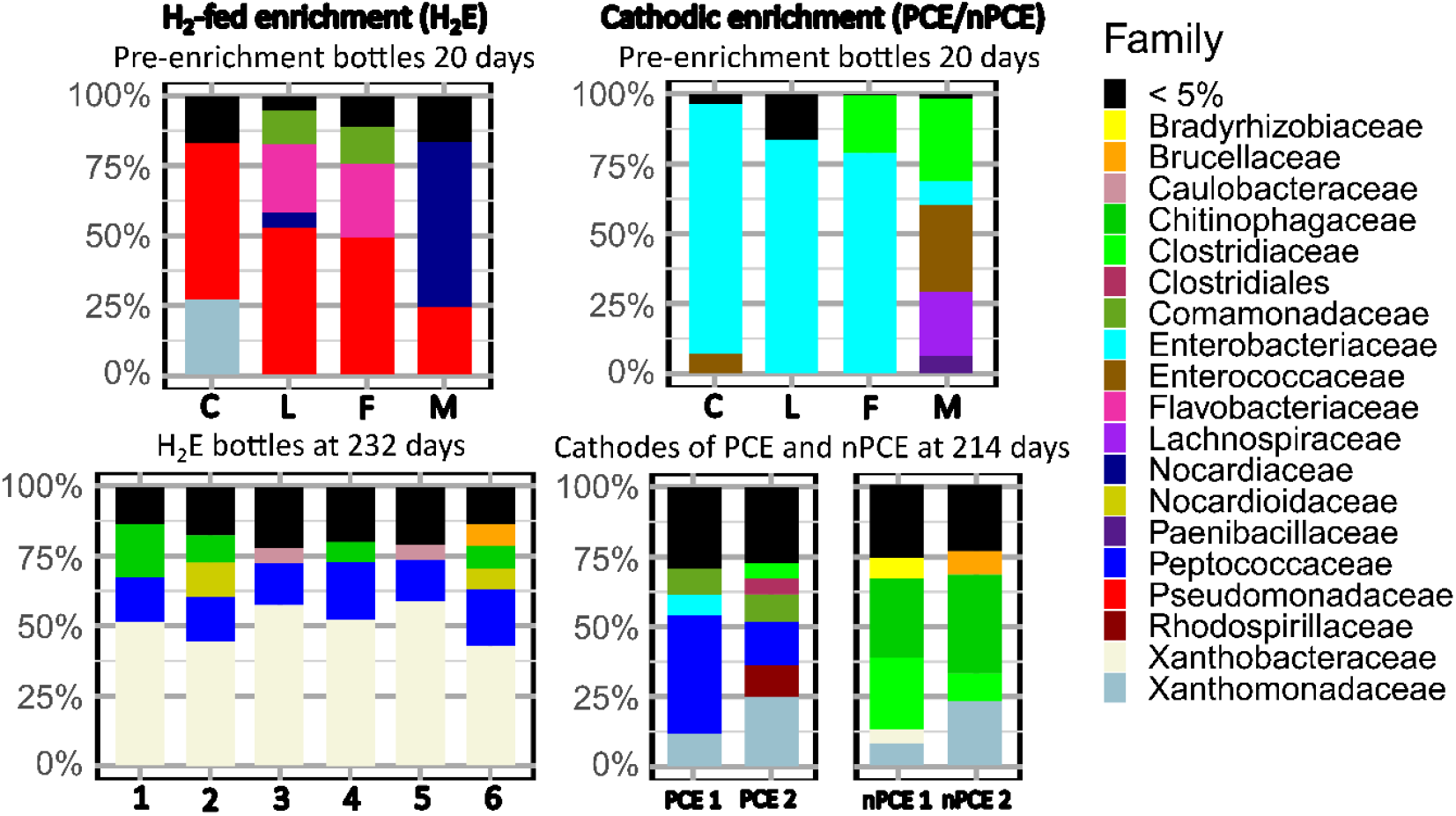
Barplots of relative abundances of major bacterial families of pre-enrichments, of H_2_-fed enrichment (H_2_E after 232 days) and cathodic enrichment (PCE and nPCE after 214 days). The microbial communities in the pre-enrichment bottles are represented by the following abbreviations: F for forest soil, C for compost, L for the rhizosphere of leguminous plants, and M for a mix of all. Only families with a relative abundance ≥ 5% are shown for each sample.

For communities at the end of the pre-enrichment, the principal component analysis (PCA) showed an important link between the families of the *Clostridiaceae, Enterobacteriaceae, Enterococcaceae* and *Lachnospiraceae* and with the PCE pre-enrichment (Figure 6a). Indeed, the group of communities of pre-enrichment with organic C is well separated from the others groups and follow the same direction as these four families. A dominance of the *Enterobacteriaceae* family (mainly of the *Citrobacter* genus) was observed for each pre-enriched sample expect for the pre-mixed sample where the three others families are highly present. These families are therefore absent or very weakly represented in the other sequenced communities as shown in Figure 7.

For the H_2_-fed bottle pre-enrichments (H_2_E), the group is also seprated of the 232-days enriched community H_2_E. *Pseudomonadaceae* (45% of average relative abundance) family was mostly dominant at the end of pre-enrichment. *Nocardiaceae, Flavobacteriales, Xanthomonadaceae* and *Comamonadaceae* were also present as seen in Figure 7. *Flavobacteriales* and *Comamonadaceae* families are also higly linked with the group of pre-enrichment in PCA of Figure 6a. These families, with the exception of *Flavobacteriales*, are also known to have members possessing the set of genes necessary for N_2_ fixation (Dos Santos et al., 2012; Ghodhbane-Gtari et al., 2019; Huda et al., 2022). These families accounted for 77% of the sequences which is high compared to the *nifH*/16S rDNA ratio of less than 0.01 at the same time point. This suggests that either the *nifH* primers were not adapted to these specific species or that the species found at this point did not possess the genes for nitrogenases. As the H_2_E pre-enrichment cultures started on a medium containing NH_4_Cl, the presence of this source of nitrogen was likely favorable to the growth of non-diazotrophic bacteria.

After 214 days of enrichment, PCE communities were affiliated to *Peptococcaceae* (29% in average), *Xanthomonadaceae* (18% in average), *Rhodospirillaceae* (11% in PCE 2, *Azospirillum*) and *Comamonadaceae* (10% in average) as presented in Figure 7. As seen in the PCA presented Figure 6b, *Rhodospirillaceae, Comamonadaceae, Enterobacteriaceae* and *Xanthomonadaceae* families are representative of the PCE cathode communities. *Peptococaccaceae* appear to be shared with communities of H_2_-fed enrichment (H_2_E) bottles. As seen in Figure 6a and Figure 7, a clear shift in microbial communities from the end of pre-enrichment was therefore observed as the difference between PCE communities at 214 days and at the end of pre-enrichment.

Only *Enterobacteriaceae* family was maintained although at minor relative abundance. Members of *Clostridiales incertae sedis* absent from original inoculum also appeared on polarized cathode. These families are known to exhibit the role of plant growth promoting bacteria (PGPB). These communities could thus be beneficial when used as living fertilizers (Cassan & García de Salamone, 2008; Rojas-Tapias et al., 2012; Singh et al., n.d.). The *Comamonadaceae* as well as the *Enterobacteriaceae* families mostly include heterotrophic species, which would be consistent with our hypotheses about the existence of interactions between heterotrophic and autotrophic populations (F. Liu et al., 2011; Wu et al., 2018). More precisely, the *Peptococcaceae* sequences were affiliated to species *Desulforamulus ruminis* (>98%). This species was already described for their ability to fix N_2_ (Postgate, 1970). The *Desulforamulus* and *Desulfotomaculum* genera have also several species able to grow with H_2_ and CO_2_ as electron and C sources (Aullo et al., 2013; Klemps et al., 1985; Zaybak et al., 2013). They were previously reported to be able to produce acetate by CO_2_ reduction through the Calvin cycle (Klemps et al., 1985), and some were already found in microbial electrochemical system on a biocathode producing acetate (Zaybak et al., 2013). The other main family, *Xanthomonadaceae*, was represented by several genera with a majority of *Pseudoxanthomonas*. In this genus, some members were identified as N_2_ fixers with a need of external organic C source, exhibiting a mixotrophic metabolism depending on the environmental conditions (J. Hu et al., 2022; Ryan et al., 2009). Sequences associated to the *Rhodospirillaceae* family were mainly affiliated to the species *Azospirillum lipoferum* which is able to grow in autotrophy with H_2_, CO_2_ and N_2_ (Tilak et al., 1986). This soil bacterium is also known for its role as a PGPB with a capacity to solubilize phosphates, making it a good candidate as a fertilizer (Cassan & García de Salamone, 2008; Tilak et al., 1986). Interestingly, many of the identified bacteria in the polarized cathode enrichment were previously described to possess the N_2_-fixing genes and capability. This supports the fact that the primers were not able to amplify the full diversity of *nifH* genes from these communities.

The *Xanthobacteraceae* (51% in average), *Peptococcaceae* (17%, identified as *Desulforamulus*), *Chitinophagaceae* (8%) and *Nocardioidaceae* (5%) families were found to be dominant in H_2_E bottles at day 214 as presented in Figure 7. The *Xanthobacteraceae* family, highly linked to H_2_E communities as seen in Figure 6b, was mostly represented by the species *Xanthobacter autotrophicus* which is known as N_2_-fixing HOB (Wiegel, 2005). This species was already been used for N_2_ fixation by Liu et al. (C. Liu et al., 2017) in an hybrid system using the H_2_ produced by a cathode. *Xanthobacter autotrophicus* was also found in the medium of the polarized cathode enrichment but in lower abundance (< 5%). Therefore, the community enriched with H_2_ in bottles was mostly composed of N_2_-fixing bacteria, as also supported by the high *nifH*/16S ratio (0.9). After 214 days of enrichment, diazotrophic HOB were specifically selected.

The presence of mixotrophic and heterotrophic bacteria in the PCE suggested that carbon-based interactions could have occurred. Acetate was the only abundant soluble carbon metabolite found in these enrichments (see Table 3). Therefore, acetate was assumed to be used as intermediate for carbon and electron transfer between autotrophic homoacetogens, e.g. *Desulforamulus rumnis*, and heterotrophic bacteria such as *Comamonas sp*.).

Furthermore the low concentration of N-NH_4_^+^ in the PCE before day 210 (Table 3) was probably due to its rapid consumption for bacteria growth. Considering these hypothesis, a conceptual scheme of microbial interactions between the main bacterial families in the PCE was proposed and is presented in Figure 8. The presence of heterotrophic bacteria and their potential use of O_2_ as a final electron acceptor was also considered. The concentration of dissolved O_2_ would have decreased in a deep layer of the biofilm due to its use by heterotrophic bacteria. A structure of the biofilm in two layers could then be proposed with a first layer composed mainly of homoacetogens fixed on the cathode and reducing CO_2_ to acetate, and a second layer composed mainly of heterotrophic bacteria using acetate and dissolved O_2_ to sustain their growth. It was assumed that bacteria in the first layer would not access to N_2_ that would be mostly fixed by organisms of the second layer.

**Figure 8.**
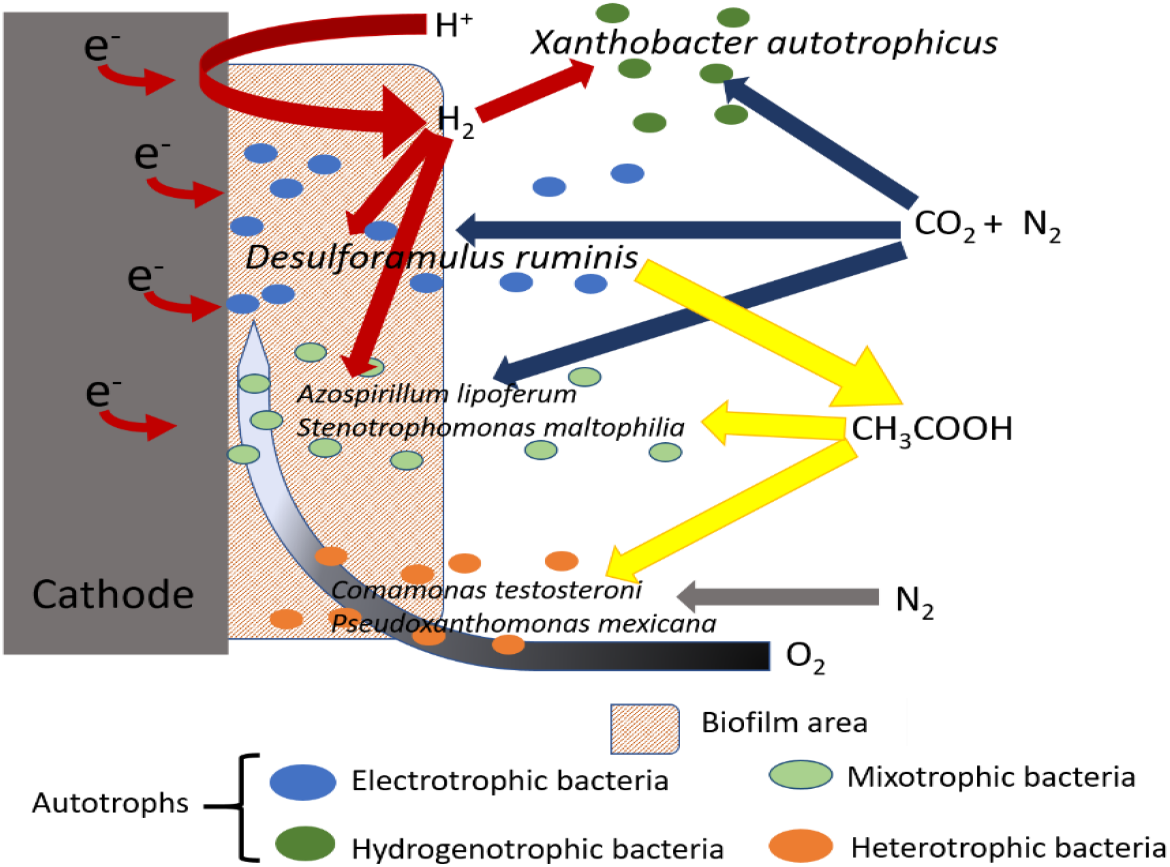
Conceptual scheme of microbial interactions occurring in polarized cathode enrichment (PCE) after enrichment for N_2_ fixation with inorganic energy and carbon sources

## Conclusion

Enrichment cultures of N_2_-fixing bacteria were successfully carried out in H_2_-fed bottles (H_2_E) and in polarized cathode enrichment (PCE). Both methods showed significant N_2_ fixation after 340 days of enrichment. The microbial communities selected were able to fix N_2_ with CO_2_ as sole carbon source and H_2_ or cathodic electrons as sole electron sources. Biomass growth on the cathode up to 4.6 10^10^ bacteria/L.d is another evidence of autotrophic growth in the PCE while bacterial growth was much lower in the H_2_E. Current density suggests the activity of autototrophic bacteria in the PCE and the availability of electron sources. As the coulombic efficiency of N_2_ fixation was low with a maximum of 3.3% and considering the low concentrations of NH_4_^+^, it was concluded that the major part of the nitrogen was incorporated into microbial biomass during the enrichment procedure. Interestingly, acetate was also produced in the PCE corresponding to a coulombic efficiency of 27%. The related microbial communities found in both enrichments had some bacterial families in common, but the communities found in the PCE appeared metabolically more diverse, suggesting probable rich microbial interactions with exchanges of electrons, carbon and nitrogen between autotrophic, heterotrophic and mixotrophic populations. Several members of the enriched communities were furthermore reported as plant growth promoting bacteria (PGPB) which could be interesting for the production of environment friendly fertilizers.To summarize, a conceptual model of microbial interactions between the main bacterial families found in the bioelectrochemical system was proposed suggesting a key role of each autotrophic, heterotrophic and mixotrophic populations in the process of N_2_ fixing by cathodic biofilms. In order to focus on the enriched microbial community and avoid to disrupt the enrichments, a comprehensive screening with different potential electron acceptors was not performed here. However, such screening would be interesting to be investigated and could be the subject of further studies on synthetic community to further explore the impact of electron acceptors on microbial communities, current densities and coulombic efficiencies.

## Acknowledgements

The authors would like to thank the INRAE Bio2E Facility (Bio2E, INRAE,2018. Environmental Biotechnology and Biorefinery Facility, https://doi.org/10.15454/1.557234103446854E12) for experimental support.

## Data and scripts availability

Data and scripts are available online on: https://doi.org/10.57745/ONNGWZ (Recherche Data gouv)

